# The sicker sex is plastic: Thermal plasticity determines sex biases in pathogen transmission

**DOI:** 10.1101/2024.11.21.624778

**Authors:** Nathan J. Butterworth, Jared Lush, Kerri J. Moore, Matthew D. Hall

**Affiliations:** School of Biological Sciences, Monash University, Clayton VIC 3800

## Abstract

Sex differences are predicted to play an important role in the spread and evolution of pathogens. However, attempts to generalize the ‘sicker’ sex have been challenged by intraspecific variability of sex-biases across the infection process. Sex specific plasticity provides a framework to resolve this by elucidating how infection is shaped at the sex-pathogen-environment interface. Using the *Daphnia magna* and *Pasteuria ramosa* system, we measure infection outcomes for males and females across three temperatures and seven pathogen densities to quantify how sex specific plasticity shapes susceptibility, pathogen loads, and ultimately transmission. We find unique forms of plasticity at each stage of infection – including equivalent, sex-specific, and divergent plasticity. Integrating these into a single estimate of transmission reveals a clear pattern – male-biased transmission at cold temperatures, and female-biased transmission at warm temperatures. Sex specific thermal plasticity thus determines the ‘sicker’ sex, with implications for pathogen spread and evolution in a warming world.

## INTRODUCTION

Throughout the animal kingdom there is often a ‘sicker’ sex (Zuk 1990; Poulin 1996; Zuk and McKean 1996; Zuk 2009; van Lunzen & Altfeld 2014; Kelly et al. 2018) defined as the sex that is most susceptible to infection (Moloo 1993; Gray 1998; Sanchez et al. 2011; Duneau et al. 2012; Morton and García-del-Pino 2013; Butterworth et al. 2024), has the highest pathogen burden (Hillegass et al. 2008; Duneau et al. 2012; Thompson et al. 2017; Gipson and Hall 2018; Rosso et al. 2020; Kailing et al. 2023), or exhibits the greatest damage from pathogen virulence (Duneau et al. 2012; Ubeda and Jansen 2016; Thompson et al. 2017; Gipson and Hall 2018; Gipson et al. 2022). Sex differences in these outcomes of infection are important because they are predicted to influence how pathogens spread and evolve (Cousineau and Alizon 2014; Ubeda and Janzen 2016; Hall and Mideo 2018; but see also Duneau and Ebert 2012, Gipson & Hall 2016 and Gipson et al. 2019). Yet attempts to identify the ‘sicker’ sex across the animal kingdom have revealed substantial variation among species (Poulin 1996; Zuk and McKean 1996; McCurdy et al. 1997; Schalk and Forbes 1997; Sheridan et al. 2000; Kelly et al. 2018) and it remains difficult to predict when and why a ‘sicker’ sex should emerge.

Attempts to generalise the identity of sicker sex have centred on the mating system of a species and the relationship between immunocompetence and reproductive effort (see Hamilton and Zuk 1982; Zuk 1990; Folstad and Karter 1992; Zuk and McKean 1996; Schwenke et al. 2016). One prediction has been that males should generally be ‘sicker’ because they can benefit from sacrificing immunocompetence and instead allocating those resources towards reproduction (Rolff 2002; Zuk et al. 2009). A challenge to this notion of such general rules, however, is emerging evidence that the ‘sicker’ sex is not static even within species. For a given species, males might be more susceptible to infection in one setting, but females more susceptible in another (e.g., Moloo et al. 1993; Gipson and Hall 2018; Gipson et al. 2022; Butterworth et al. 2024). While it is well established that host-pathogen interactions are highly sensitive to environmental context (Gervasi et al. 2015; Gehman et al. 2018; Mordecai et al. 2019; Kirk et al. 2019; Hector et al. 2024),d that the environment can differentially modify the characteristics of disease in males and females (i.e., ‘sex-specific plasticity’) has received scant attention. Considering the widespread importance of sex-specific plasticity in response to changing environments (as per Bonduriansky 2007; Stillwell et al. 2010; Stillwell and Davidowitz 2010; Rohner et al. 2017; Pottier et al. 2021; Hangartner et al. 2022; Brand et al. 2023; Polverino et al. 2023), uniting sex-specific plasticity with our understanding of disease provides an opportunity to resolve and improve our understanding of the ultimate rules shaping the ‘sicker’ sex across the animal kingdom.

Opportunities for sex-specific plasticity to influence disease dynamics can occur multiple times during any encounter between a host and pathogen (sensu Hall et al. 2017), from the initial attempt by the pathogen to establish in the host (susceptibility), though to its proliferation within the host (pathogen load), and any damage caused (virulence). At each of these stages of infection, however, different forms of sex-specific plasticity can emerge. For example, a given component of infection could be (1) plastic only (or to a greater extent) in one sex (i.e., sex-specific plasticity, Figure 1A), (2) plastic in both sexes in the same direction (i.e., equivalent plasticity, Figure 1B), or (3) plastic in both sexes but in opposing directions (i.e., divergent plasticity, Figure 1C). Because the extent and form of plasticity can be distinct for each stage of infection – males may be ‘sicker’ *and* more plastic in one trait (i.e., susceptibility), while females are ‘sicker’ *but less* plastic in another (i.e., pathogen burden) – hampering attempts at a simple conclusion to which is the ‘sicker’ sex. However, models of infectious disease provide a way forward by summarising outcomes across the stages of infection into a single metric of pathogen performance which is comparable among systems – pathogen transmission (Anderson and May 1979; de Jong et al. 1995; McCallum 2001; Grassly and Fraser 2008). In other words, we can use models of disease transmission to redefine the ‘sicker’ sex simply as the one most likely to transmit the pathogen to other individuals (i.e., ‘per-capita transmission’, Figure 1D) – which provides a clear way to interpret the overall consequences of sex-specific plasticity for disease dynamics (Figure 1).

**Figure 1.**
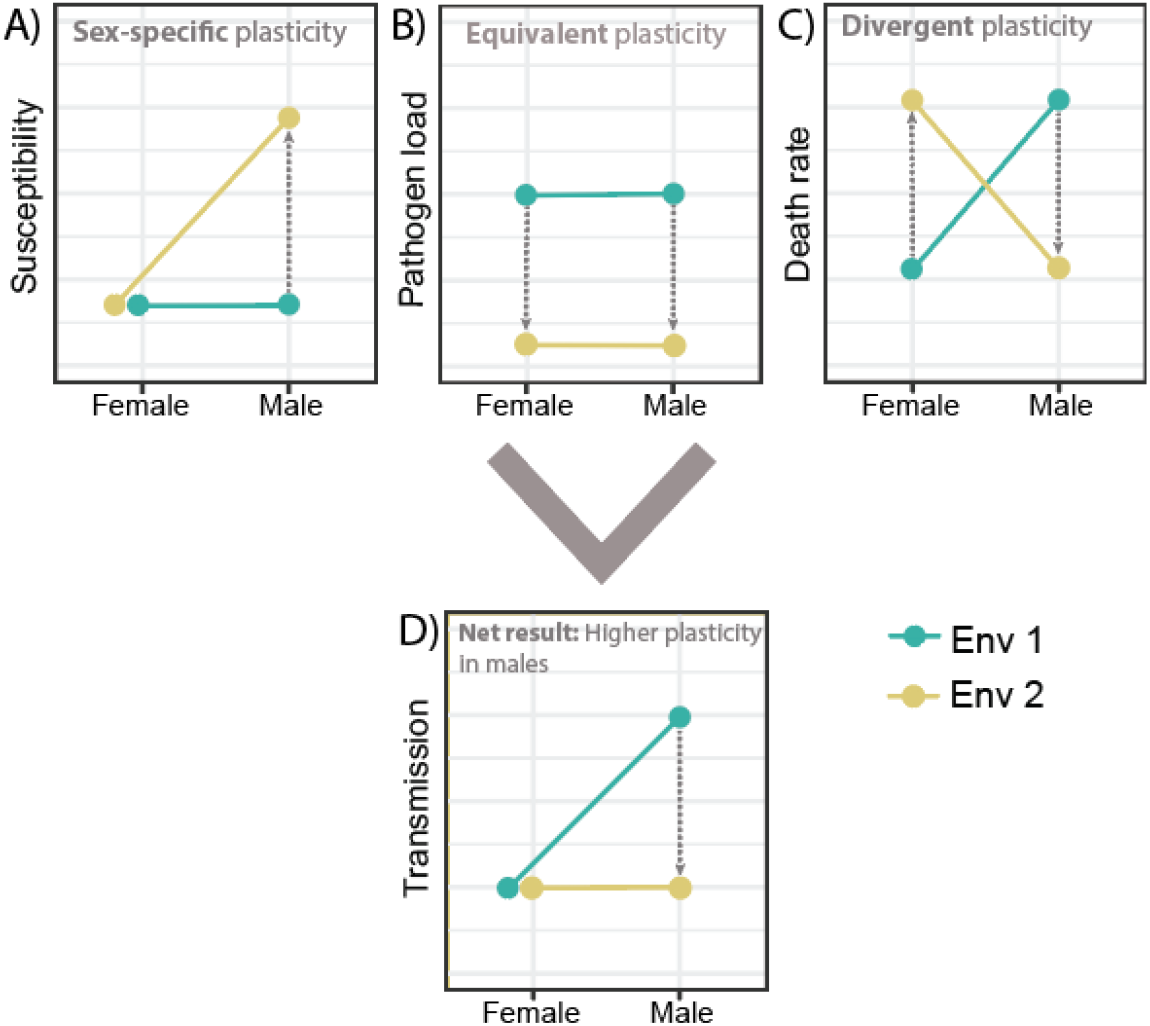
Different forms of sex-specific plasticity in each component of infection can lead to non-intuitive predictions for sex-biases in pathogen transmission. Motivated Using a horizontally transmitted pathogen as an example (e.g., a bacterial infection of *Daphnia magna* that is transmitted only at host death) the capacity for pathogen transmission through each sex will depend on the sex-specific differences in host susceptibility, pathogen loads, and host death rates (e.g., Hall and Mideo 2018; Aulsebrook et al. 2023). Hypothetically, each component of infection could have a unique form of sex-specific plasticity – susceptibility might be plastic only for males (i.e., sex-specific plasticity), pathogen loads equally plastic in both sexes in the same direction (i.e., equivalent plasticity), and death rates equally plastic in both sexes but in opposing directions (i.e., divergent plasticity). Yet non-intuitively, integration each component into a single measure of disease spread would suggest a strong male-bias in transmission in environment 1, but no sex bias at all in environment 2, and that this shift in the extent of the ‘sicker sex’ (in regard to the sex where pathogen transmission is highest) would be underpinned entirely by plasticity in male transmission only.

Of all environmental factors, temperature provides an ideal opportunity to assess the contribution of sex-specific plasticity to sex-biases in disease transmission. For one, understanding the plastic responses of both sexes and their pathogens to temperature is increasingly important, as climate change is expected to fundamentally alter the spread of disease (Gehman et al. 2018; Mordecai et al. 2019). Thermal plasticity has been shown to occur across every step of the infection process – from a host’s susceptibility to pathogen growth rates, and virulence (Gehman et al. 2018; Kirk et al. 2019; Mordecai et al. 2019; Shocket et al. 2019; Hector et al. 2024) with contrasting forms of plasticity arising at each component (Kirk et al. 2018; Kirk et al. 2019; Hector et al. 2024). Sex-specific responses to temperature have been observed across the animal kingdom for body size (Stillwell et al. 2009; Stillwell and Davidowitz 2010), movement activity (Brand et al. 2023) and thermal tolerance (Pottier et al. 2021; Hangartner et al. 2022). Surprisingly, although well known that males and females widely exhibit differential plastic responses to temperature – how sex-specific plasticity and the thermal dependency of disease combine to shape sex differences in the transmission of pathogens remains unknown.

To understand how sex-biases in transmission emerge, and the role that the thermal environment might play, a crucial consideration is the density of pathogen propagules encountered – which determines the likelihood and consequences of contact between the host and pathogen (Regoes et al. 2002; Regoes et al. 2003; Ben-Ami et al. 2010; McCallum et al. 2017; Clay et al. 2021) and can shape all subsequent components of infection (Ebert et al. 2000; Regoes et al. 2002; Regoes et al. 2003; Ben-Ami et al. 2010; McCallum et al. 2017; Lunn et al. 2019; Clay et al. 2021). Put simply, at low doses, encounters with the pathogen are rare and immune systems remain unchallenged, providing little opportunity for sex differences in immunity, for example, to be expressed. Conversely, at high doses pathogen encounter rates are maximised and immune systems easily overwhelmed, even for the most resistant sex. Both scenarios have been shown to obscure sex-biases in susceptibility (Butterworth et al. 2024) and to alter the sensitivity of different components of infection to environmental change (e.g., Kirk et al. 2019). Considering the dose-dependency of thermal plasticity across the entire infection process thus expands the scope for sex-specific plasticity to emerge or to be constrained.

To investigate how sex-specific thermal plasticity emerges across the infection process and shapes sex-biases in pathogen transmission, we use the *Daphnia magna* and *Pasteuria ramosa* system (Ebert et al. 2016) to measure infection outcomes across three temperatures, seven pathogen densities, and 2,100 individual clonal males and females. Previous studies have shown that thermal plasticity is sensitive to initial pathogen dose (Kirk et al. 2019) and fundamentally determines infection rates in *Daphnia* (Kirk et al. 2019; Shocket et al. 2019). Males and females also exhibit strong sexual dimorphism across almost every measurable trait – males being significantly smaller (Duneau et al. 2012), having lower thermal tolerances when uninfected (Laidlaw et al. 2020), differing in susceptibility to infection (Butterworth et al. 2024), and with lower pathogen burdens (Gipson et al. 2022). Considering the thermal sensitivity of the infection process, strongly sexually dimorphic infection outcomes, and differing thermal sensitivity between the sexes – the *Daphnia*-*Pasteuria* system provides the ideal opportunity to explore how sex-specific plasticity can shape the extent and identity of the ‘sicker’ sex.

## METHODS

### Study organisms

The freshwater crustacean *Daphnia magna* Straus (Cladocera: Anomopoda) reproduces via facultative parthenogenesis and can produce genetically identical male and female clones. In its natural environment it is frequently parasitised by a range of organisms (Ebert et al. 2005), including the gram-positive bacteria *Pasteuria ramosa* Metchnikoff (Green 1974), which attaches to the host oesophagus and reproduces within the haemolymph to produce transmission spores (Ebert et al. 2016). Infected females suffer severely reduced fecundity and lifespan, and an increase in body size (Clerc et al. 2015; Hall et al. 2019). Males suffer less with fewer spores and a smaller reduction in longevity (Duneau et al. 2012; Gipson et al. 2022). Transmission is exclusively horizontal, only occurring after the release of spores from dead hosts (Hall and Mideo 2018; Aulsebrook et al. 2023).

### Production of experimental animals

This experiment used stocks of *D. magna* of the genotype BE-OHZ-M10 which was originally hatched from a resting egg (ephippia) collected from Oud Heverlee, Belgium (coordinates: 50° 50′ 14”, 4° 39′ 48′′). The *P. ramosa* genotype C1 was used to infect which was originally isolated from a single female infected with pond sediment from Moscow, Russia (Luijckx et al. 2010). This host-pathogen combination was chosen because they are compatible and have been shown to result in sex-biases in pathogen loads (Laidlaw et al. 2020). To minimise variation in maternal effects, standardised female *D. magna* were raised individually for three generations in 70 mL jars filled with 50 mL of artificial *Daphnia* medium (ADaM – Klüttgen et al. 1994; modified by Ebert et al. 1998) and were maintained under standard conditions (20°C, 16L:8D).

Genetically identical male and female *D. magna* were produced by exposing the third generation of standardised females to the crustacean juvenile hormone methyl farnesoate (300ug L; product ID: S-0153, Echelon Biosciences, Salt Lake City, UT). Following established protocols, adult females were transferred to 20 mL of hormone after the release of their first clutch (Thompson et al. 2017; Gipson et al. 2022). The second clutch produced after exposure to the hormone, composed of a mixture of male and female offspring, was then collected and immediately returned to normal artificial media and placed into the relevant temperature treatment (15, 20, or 25°C). This method of producing both male and female *Daphnia* has no detectable impact on either the lifespan or fecundity of control animals (Thompson et al. 2017). Male and female *Daphnia* offspring were identified by the presence/absence of the sexually dimorphic appendage used for clasping onto females (Ebert 2005).

### Experimental design

To assess how the sensitivity of thermal plasticity across environmental doses, we performed a dose-response experiment across three temperatures. Males and females to be infected were housed individually at their respective temperature (15, 20, or 25 °C), in 70 ml jars, filled with 50 ml of artificial media. Animals were fed daily throughout the entire experiment with increasing numbers of algal cells (Scenedesmus sp.) – 1 million cells (at 1 day old), 2.5 million cells (at 2-5 days old), 3 million cells (at days 5-6 days old), and 5 million cells (from day 7 onwards). Animals were changed into 50 ml of fresh ADaM every three days throughout the entire experiment. Each treatment consisted of 50 individual replicates for a total of 2,100 experimental animals (2 sexes × 3 temperatures × 7 doses × 50 replicates).

To infect the animals, *P. ramosa* spores were set up as follows: the highest spore dose of 1,250,000 spores was subsequently diluted seven times by a factor of five to produce the other doses, resulting in seven dose levels of 80, 400, 2000, 10,000, 50,000, 250,000, and 1,250,000 spores. At four days of age, the animals were transferred individually into jars containing 20 ml of ADaM and dosed with one of the seven *P. ramosa* spore doses and immediately returned to their respective temperature cabinet. After 4 days, animals were moved to fresh jars filled with 50 ml of ADaM and then water changed as usual after.

### Quantifying infection rates, pathogen loads, death rates, and virulence

Experimental animals were monitored daily throughout the experiment until death at which point their lifespan was recorded, and they were frozen in 500 uL of purified water for later determination of infection status and measurement of transmission spores. All *Daphnia* were thawed, crushed with a pestle, and sampled individually for infection status and spore count by haemocytometer counts under a phase-contrast microscope. Death rate was calculated as the inverse of lifespan (1/lifespan). Virulence was calculated as the difference between the mean lifespan of uninfected animals taken from the lowest dose (80 spores) and a given infected individual. The experiment started with 2100 animals, and 2004 were successfully assayed. 96 animals were removed due to being lost or mislabelled during the experiment. All following statistical analyses were performed in R (ver. 4.0.2; R Development Core Team 2023).

### Assessing the sex-specific plasticity of susceptibility

To test how sex biases in susceptibility to infection emerge across temperatures, we first modelled the change in the proportion hosts infected with increasing spore dose as a three-parameter logistic non-linear model via the package *drc* (where *Y* = *d*/(1 + *e*^*-b(X-e)*^); Ritz et al. 2016), allowing the partitioning of susceptibility into three components: e, the relative ED50 (the dose at which 50% of susceptible hosts are infected), b, the slope (related to the variance in susceptibility), and *d*, the max asymptote (the maximum proportion of infected hosts). For each temperature, we compared the fit of a model where the parameters were constrained to be equal between the sexes to one where we allowed the parameters to vary by sex (as per Butterworth et al. 2024). At the four doses with appreciable infection rates (10,000, 50,000, 250,000 and 1,250,000 spores), we also analysed the probability of infection using a binomial generalised linear model with a logit link function from ‘glm’ function using the ‘stats’ package (v 4.2.2) with sex, temperature, and dose as fixed effects.

### Analysing the plasticity of sex-biases across the entire infection process

Next, at the four doses with appreciable infection rates (10,000, 50,000, 250,000 and 1,250,000 spores) we assessed how sex-biases in host lifespan, virulence, and total spore loads varied among dose, sex, and temperature treatments. For death rates we used square root transformation and a linear model with sex, temperature, and dose as fixed effects. For virulence, we used a linear model with the ‘lm’ function from the ‘stats’ package with sex, temperature and dose specified as fixed effects. For spore loads, we used a square root transformation and a generalised least squares model with sex, temperature and dose as fixed effects using the ‘gls’ function from the ‘nlme’ package (v. 3.1, Pinheiro et al. 2023) and allowed for residual variance to vary independently for each temperature treatment to account for heteroscedasticity using ‘VarIdent’. We then used ‘Anova’ type III from the ‘car’ package (v. 3.1-1, Fox and Weisberg 2019). Predicted mean infection rates, spore loads, death rates, and virulence and standard errors were extracted from each model with ‘emmean’ function from the ‘emmeans’ package (v. 1.8.3, Lenth et al. 2024).

### Quantifying sex biases in pathogen transmission

To understand how sex biases in pathogen transmission were mediated by sex-specific plasticity, we use the composite transmission rate (σ = β□ωD) adapted from Hall and Mideo (2018) and following Aulsebrook et al. (2023), which describes the per capita rate at which new infections are created based only on the characteristics of infected individuals. This term will increase with an increase in environment transmission rates (β), spore loads at host death (ω) or the rate at which infections end owing to host mortality (D). It also depends on the effective production rate of free-living environmental spores (□).

To calculate transmission (σ) for each temperature, dose, and sex combination we quantified the relevant components using data from the experimental animals that were observed from birth until death. For transmission, spore loads at host death (ω) were available for each infected individual and we estimated the rate at which infections end owing to host death (D) as the inverse of measured host lifespan (1/days). The environmental transmission rate (β) was estimated using the numbers of infected and uninfected individuals from each treatment, where the probability of remaining uninfected (P) depends on the density of pathogen spores (Z, Dose per 20 ml) and the length of the infection period (t, 4 days), such that P = e^−βZt^ (following Shocket et al. 2018).

For all traits we used the Stan modelling language (Stan Development Team 2023) as implemented via the CmdStanR package v. 0.5.3 (Gabry et al. 2023) in R, to calculate Bayesian posterior distribution estimates for each metric (see also Shocket et al. 2018). We used the default sampling settings of Stan and assumed uniform priors based on a plausible range of values (following Aulsebrook et al. 2023). In all following analyses, we arbitrarily set = = 0.01 (following Hall and Mideo 2018) as it was not measured in this experiment. To calculate the composite transmission function (σ) we fed the posterior samples of each of the metrics described above into the composite transmission rate equation, and then characterized the resulting distribution via its mean and 90% credible intervals (capturing that 95% of the mass in this interval is above or below the outer values).

## RESULTS

### The more susceptible sex arises through sex-specific thermal plasticity

To characterise how sex biases in susceptibility are mitigated by the relationship between environmental temperature and pathogen dose we first estimated each dose-response curve for the probability of becoming infected using a three-parameter log-logistic model. At 15 °C, we found that males were broadly more susceptible to infection (Figure 2A), as allowing parameters to vary by sex significantly improved the fit of the dose-response model (F_3,8_ = 8.468, *p* = 0.007). At 20 °C we found no statistical support for differences between males and females in their dose-response curves (F_3,8_ = 0.711, *p* = 0.572). Thus, sex differences in susceptibility did not appear to arise at 20 °C (Figure 2A). However, at 25 °C we again found that allowing parameters to vary by sex significantly improved the fit of the dose-response model (F_3,8_ = 50.091, *p* < 0.001). Yet the opposite trend was observed relative to the 15 °C treatment, with females now more susceptible to infection across most doses (Figure 2A).

**Figure 2.**
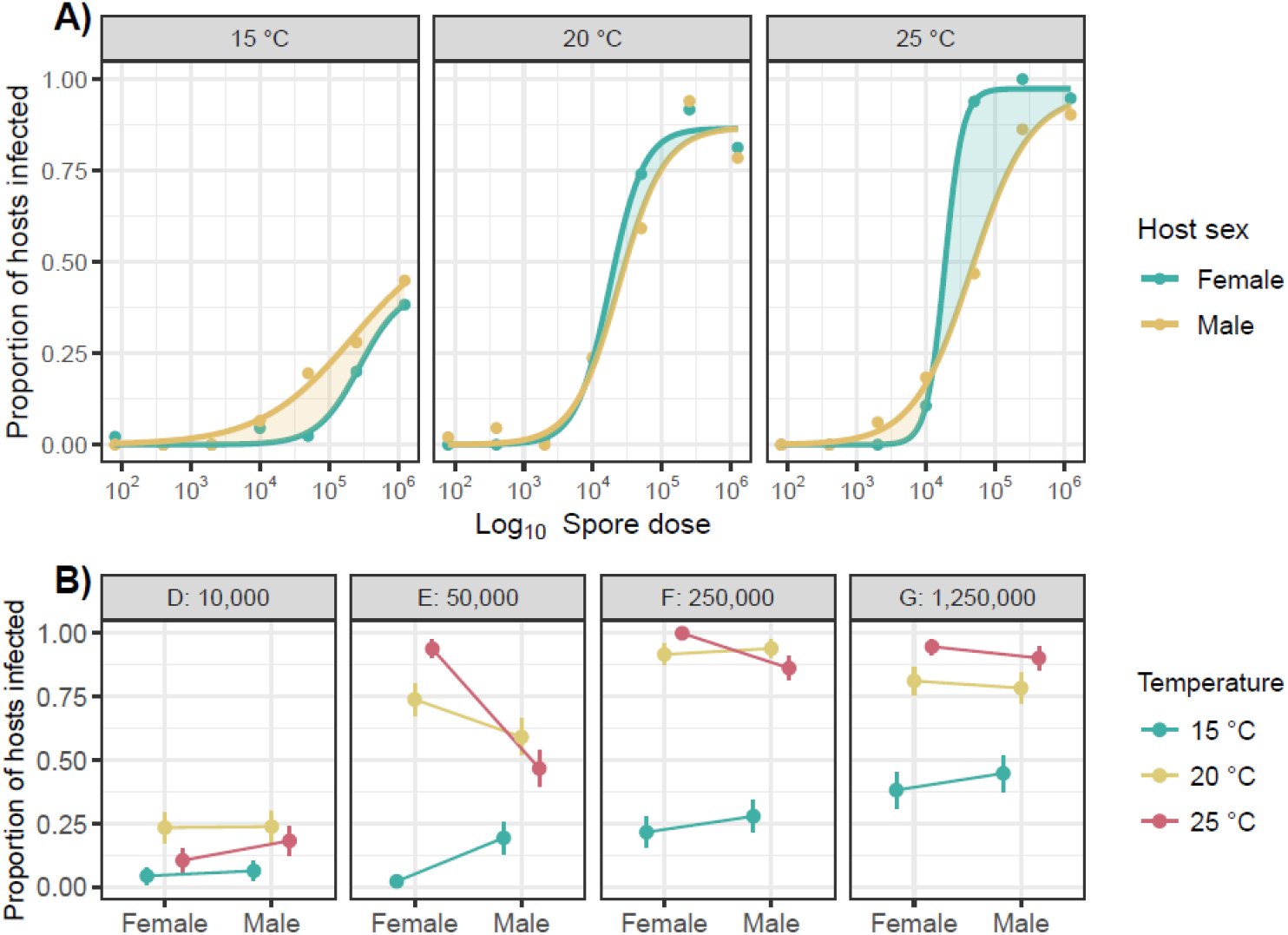
A) The nonlinear relationship between pathogen dose and infection success. Shown is the change in proportion of infected *Daphnia magna* (genotype: BE-OHZ-M10) hosts in relation to the dose of the pathogen *Pasteuria ramosa* (genotype: C1), as modelled by a three-parameter logistic model with log_10_ spore dose as the explanatory variable. Hosts were exposed to the pathogen when four days old and at one of three temperatures (15 °C, 20 °C, 25 °C). Yellow shading represents a male bias in the probability of infection. Green shading represents a female bias. B) Predicted means and standard errors of infection rates from the binomial generalised linear model at each of the four doses where infection rates were appreciable.

To further interrogate how thermal plasticity in sex biases arises at each dose, we next analysed only the doses where rates of infection were appreciable (>10,000 spores, Figure 2B). After doing so, we found infection rates depended on a three-way interaction between dose, temperature, and sex (Table 1). Thus, while temperature directly mediates the magnitude and direction of sex biases in infection rates, the pattern was not consistent across doses. At the lowest and highest doses there were no sex biases in susceptibility and only an effect of temperature on overall infection likelihood (Figure 2B, Table 1). Instead, sex biases emerged at the intermediate doses (50,000 and 250,000) and this is where the effect of temperature was felt most strongly on plasticity in sex biases. As expected by the dose-response curves (Figure 2A), the 15 °C treatment favoured higher infection rates in males, while the 25 °C favoured higher infection rates in females, leading to a two-way interaction between sex and temperature at each dose (Table 1).

**Table 1.**
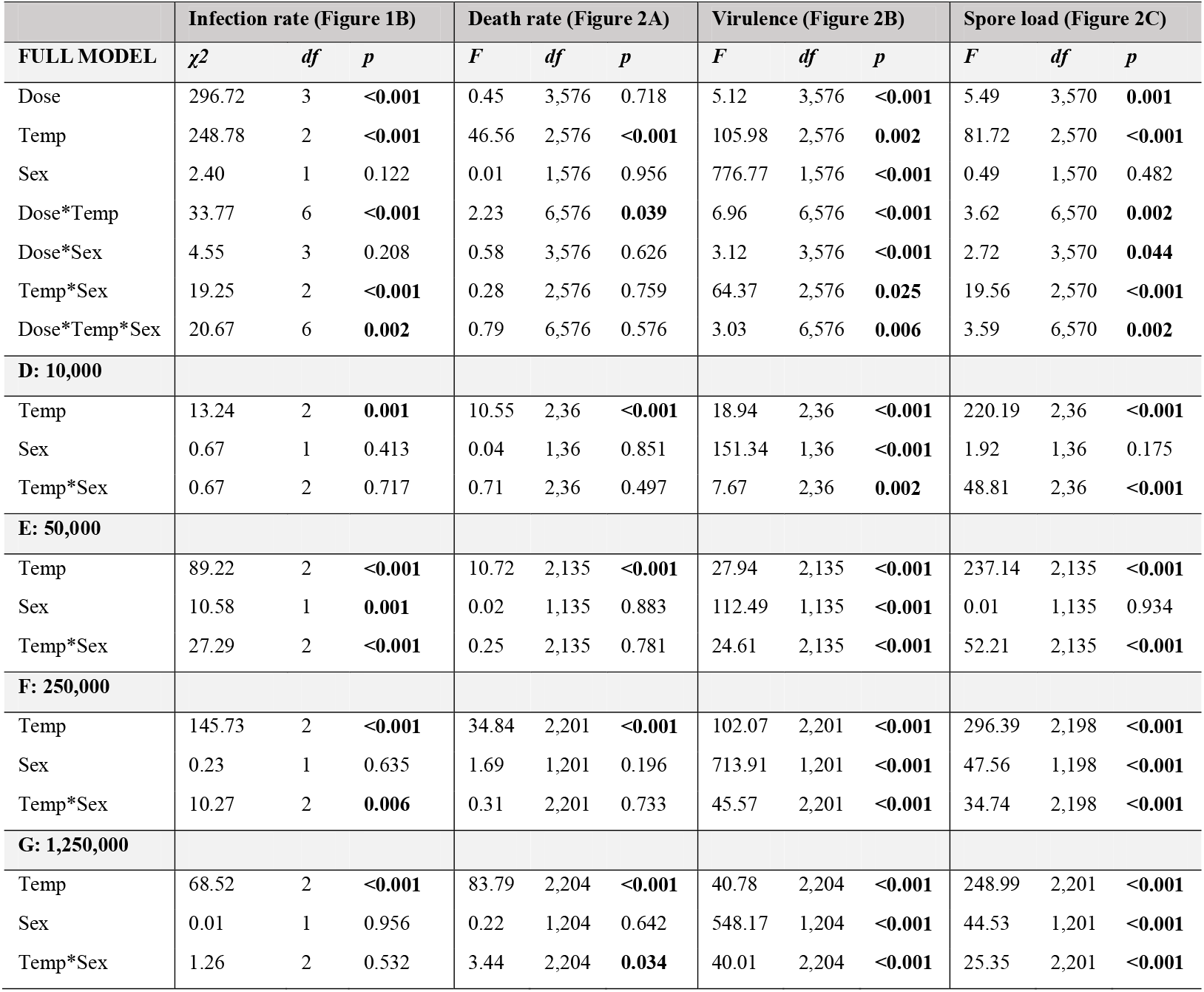
Result of the analysis of variance (type III) for the effects of dose (4 levels: 10,000, 50,000, 250,000, and 1,250,000 spores), sex (2 levels: male or female), and temperature (3 levels: 15 °C, 20 °C, and 25 °C) on infection prevalence (binomial generalised linear model on the probability of being infected), death rate (linear model on square root death rate), virulence (linear model on the reduction in lifespan relative to the control average), and spore load (generalised least squares on the square root total number of spores at host death). Bold numbers indicate significant values.

### Patterns of plasticity were not reinforced in downstream components of infection

Downstream of infection, the effects of temperature on sex-biases were not felt equally for subsequent infection outcomes – with each component showing a unique functional form of plasticity. For the death rates of infected animals, there was no overall sex bias with death rates simply increasing with temperature (Figure 3A). Broadly then, death rate shows equivalent thermal plasticity in both sexes (i.e., no overall sex-by-temperature interaction, Table 1). In contrast, spore loads at host death showed clear differences in the thermal plasticity of each sex, and the degree of sex biases occurring at each temperature as a result (Figure 3B, and sex-by-temperature interactions in Table 1). Females increase by fifteen-fold, especially at low doses. Males, in contrast are much less sensitive increasing only four-fold. As a result, sex biases in pathogen load are driven by greater thermal plasticity in females. Overall, the significant three-way interaction between dose, temperature, and sex-biases reflects the subtle effect that pathogen dose has on the interaction between temperature and sex (Table 1).

**Figure 3.**
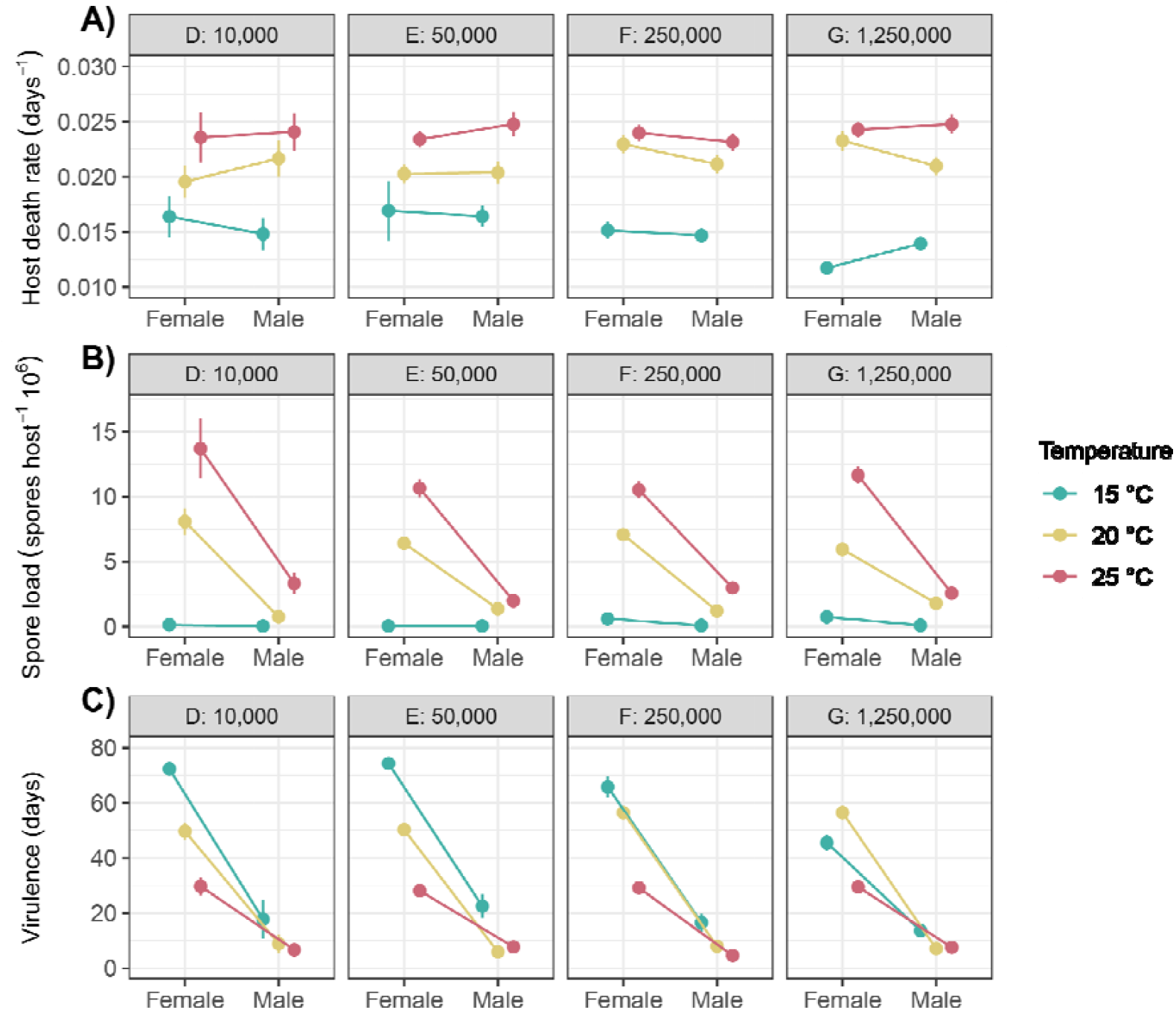
Thermal plasticity across components of the infection process as measured for *Daphnia magna* (genotype: BE-OHZ-M10) exposed to the pathogen *Pasteuria ramosa* (genotype: C1). Shown are the predicted treatment means (± SE) for A) host death rate, B) host virulence (reduction in lifespan relative to uninfected hosts), and C) the number of mature spores present within the host upon death. Each facet represents one of the four exposure doses where infection rates were appreciable as per Figure 2A.

Mirroring the observed variation is spore loads was the reduction in host lifespan caused by the pathogen (i.e., virulence, Figure 3C). Although the death rates of infected animals did not differ between the sexes (see above and Figure 3A), the lifespan of control males and females did at each temperature (see supp. material 1, Figure S1). This allowed for sex biases in virulence to emerge, which again were due to greater female sensitivity to temperature (Figure 3C). Accounting for the significant three-way interaction between dose, temperature, and sex-biases there was a rank order change across doses in the temperature with the greatest reduction in female lifespan – which was 15 °C at low doses, but 20 °C at the highest dose (Table 1).

### Warmer temperatures increase sex-biases in transmission through greater female plasticity

Integrating each characteristic of infection into a single measure of pathogen performance, reveals that sex biases in pathogen transmission depend on the interaction between dose and temperature (Figure 4A). Warmer temperatures led to higher tranmission in general, with females most plastic in their response temperature at every dose (Figure 4A), leading up to a 48-fold increase towards female-biased transmission at intermediate doses. This result reflects the higher thermal plasticity females showed in key components of pathogen transmission – infection rates (Figure 2B) and spore loads at host death (Figure 3B). However, transmission was not exclusively female biased. At low temperatures (15 °C), there was up to a 13-fold shift towards male-biased tranmission at lower doses (particularly at the 50,000 spores/ml dose) and a lack of dimorphism across other doses (Figure 4B). Overall, the most pronounced manifestation of sex differences occurs at dose E (50,000 spores) across all temperatures.

**Figure 4.**
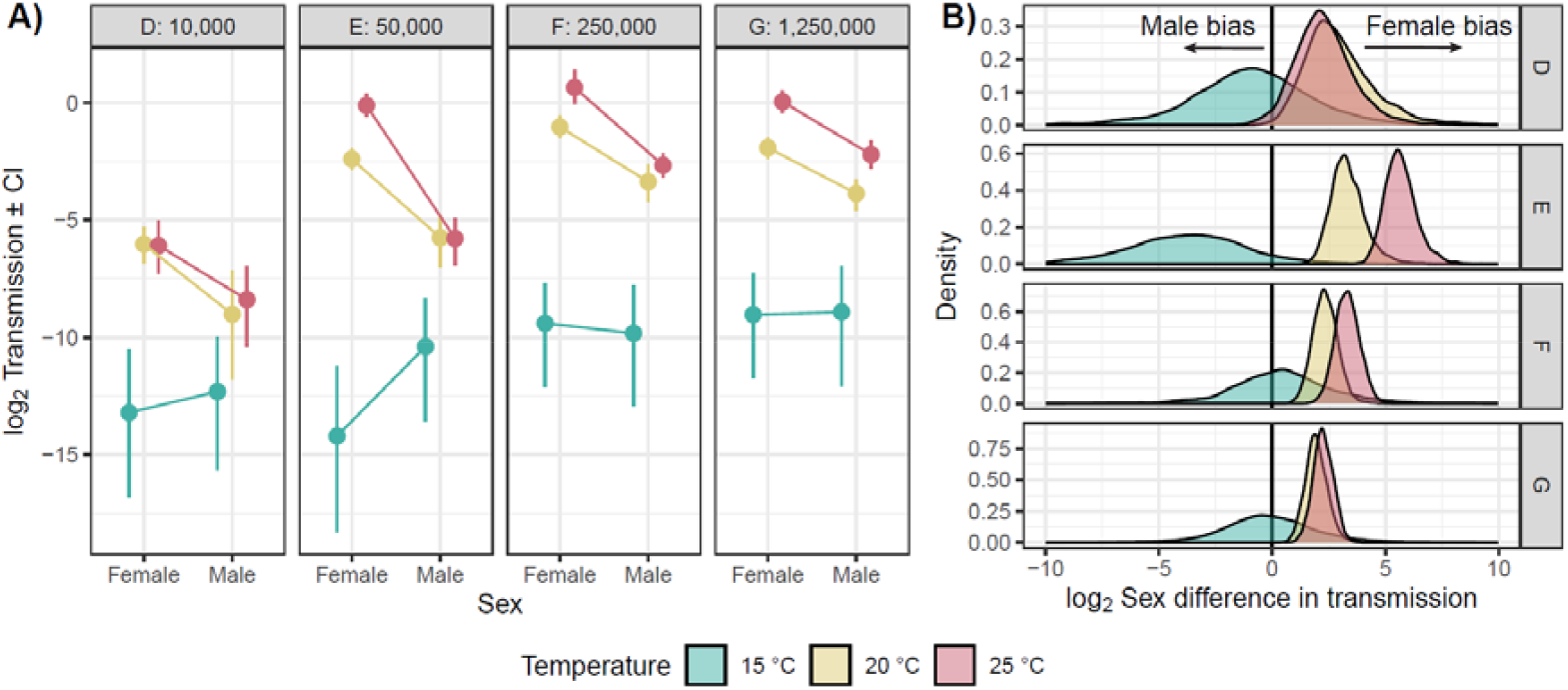
Thermal plasticity in transmission (calculated as the composite transmission rate from Hall and Mideo 2018; Aulsebrook et al. 2023) as displayed on the log_2_ scale. Shown are A) the treatment means and 90% credible intervals for the four doses that had appreciable infection rates (as per Figure 2A), and B) the extent of sex difference in transmission calculated as female value minus the male value for each of the four doses that had appreciable infection rates (as per Figure 2A).

## DISCUSSION

The extent of sex-biases in transmission are predicted to fundamentally shape the spread and evolution of pathogens in species with pronounced sexual dimorphism (Cousineau and Alizon 2014; Ubeda and Jansen 2016; Hall and Mideo 2018). A hallmark of male-female differences for many species, however, is their differential plasticity in response to environmental change (Bonduriansky 2007; Stillwell et al. 2010; Stillwell and Davidowitz 2010; Rohner et al. 2017; Pottier et al. 2021; Hangartner et al. 2022). Here, we integrate sex-specific plasticity with an understanding of sexual dimorphism in pathogen transmission. By manipulating the thermal environment, we show that temperature fundamentally determines the extent of sex-biases in many metrics of infection, and ultimately transmission. The identity of the ‘sicker’ sex thus depends not only on the species in question (Poulin 1996; Zuk and McKean 1996; McCurdy et al. 1997; Schalk and Forbes 1997; Sheridan et al. 2000) but is also determined by the environmental context, and likely highly variable even within-species. While our results challenge the notion of a static ‘sicker’ sex within species, they also highlight the remarkable flexibility of the sicker sex across environments and broaden the scope in which we can search for evolutionary generalizations of sex biases in infection outcomes.

In this single host-pathogen system we saw every possible form of sex-specific plasticity that could arise (e.g., Fig 1). For some components of infection, such as death rate, there was no sex bias and each sex responded equally to thermal change (e.g., Fig 1B) leading to a general rise in death rates with warmer temperatures (see Fig 3A). For other components, plasticity was sex-specific (e.g., Fig 1B), with females far more sensitive to temperature changes than males. In the case of pathogen load, rising temperature drove ever increasing difference between the sexes (Fig 3B), while the reverse pattern was observed for the reduction in lifespan (Fig 3C). Finally, for infection rates, we found evidence for divergent sex-specific plasticity (e.g., Fig 1C), as males had generally higher infection rates at 15°C and females higher infection rates at 25°C (Fig 2A), although these patterns were conditional on initial pathogen dose (Fig. 2B). These results highlight that the very definition of the sicker sex is both entirely trait dependent and highly modifiable by environmental change (see also Gipson et al. 2022; Butterworth et al. 2024). An absence of a sex bias under one environmental condition thus does not preclude the emergence of a male or female bias in another.

Integrating the different components of infection into a single measure of transmission, derived from formal epidemiological models (Hall and Mideo 2018; Aulsebrook et al. 2023), redefined the sicker sex as the one most likely to spread disease. In doing so, it became evident that warmer temperatures favoured increasingly female-biased pathogen transmission, driven by the greater sensitivity of females to warming (Fig 4). Only at the coldest temperature, and the lowest initial doses, did we observe a shift towards male-biased transmission (Fig 4). In terms of the spread of infectious disease, therefore, the brunt of rising temperatures will most heavily be felt though females in this system (see also Gipson et al. 2022 for a similar argument regarding resource fluctuations). Indeed, females suffering the greatest consequences of warming on traits underlying pathogen transmission could be a widespread trend. Although neither males nor females are universally more responsive to warming on average (i.e., Pottier et al. 2021, Hangartner et al. 2022), for a subset of species, females have been shown to be the more thermally plastic sex for ecologically relevant functions such as thermal tolerances (e.g., for five of 11 *Drosophila* species, Mitchell& Hoffmann, 2010) and growth rates (Stillwell et al. 2010). However, there are also many examples where males exhibit greater plasticity to temperature in their thermal tolerance (e.g., two of 11 *Drosophila* species, Mitchell& Hoffmann, 2010) or other fitness components (e.g., body size, Stillwell and Fox 2007; locomotor activity, Brand et al. 2023). Thus, in other systems where males are the more thermally plastic sex, we may see a shift towards male-biased pathogen transmission as temperatures rise.

Why might female *D. magna* show greater thermal plasticity in infection outcomes? Thermal plasticity has been shown to scale with body size across species (see Rohr et al. 2018), with a greater capacity for phenotypic plasticity occurring for the larger sex in several other invertebrate species with strong sexual dimorphism (Teder and Tamaru 2005; Bonduriansky 2007; Stillwell et al. 2010; Rohner et al. 2018). In *D. magna*, females are significantly larger than males (Duneau et al. 2012), suggesting a greater capacity for plasticity already (sensu Rohr et al. 2018), but they also display higher and more environmentally sensitive feeding rates (e.g., Gipson et al. 2022). Because feeding rate is linked directly to pathogen contact rate in this system (Hall et al. 2007) and warming directly increases feeding rates (Shocket et al. 2018; Shocket et al. 2019), it is likely that females intake disproportionately more resources and pathogens compared to males as temperatures rise. Thus, thermal sensitivity is higher for both female infection rates (likely due to greater increases in pathogen contact rates with warming for females, Hall et al. 2007; Shocket et al. 2019) and spore loads (likely due to greater increases in available energy for pathogens with warming in females; Nørgaard et al. 2021).

That the direction and magnitude of sex differences in pathogen transmission depends so strongly on the environmental context also has important consequences for the evolutionary trajectories of pathogens. When pathogens can encounter and infect multiple types of hosts (i.e., host heterogeneity), theory suggests pathogens will become strongly adapted to the host type which they most frequently encounter and through which they have the highest transmission (see Gandon 2004; Osnas and Dobson 2012). This applies equally to sex differences, which are just another form of host heterogeneity (Cousineau and Alizon 2014; Úbeda and Jansen 2016; see also Duneau and Ebert 2012; Gipson and Hall 2016). For example, when it is warm and transmission is maximised predominantly in females, such as observed here in *Daphnia*, females are predicted to have a greater impact on pathogen evolution, with any impact of host sex more evenly shared under cooler conditions (or even slightly male-biased). In *Daphnia*, at least, females are expected to favour more virulent pathogen genotypes (Gipson et al. 2019), which our results suggest would likely lead to a rise in virulence under warming. But for other systems the evolution of overall virulence will depend on the magnitude of between-sex transmission and the relative timing of propagule production and mortality during infection (Cousineau and Alizon 2014; Úbeda and Jansen 2016). Thermal changes can thus shape the sex that drives pathogen adaptation, with significant implications for the evolution of pathogen virulence – an aspect often overlooked in the context of global change.

In conclusion, we have revealed that the identity of the “sicker” sex within a species is not fixed, but rather contingent upon the environment and the patterns of sex-specific plasticity that emerge at each stage of the infection process. As global temperatures continue to rise, our results show how sex-specific plasticity can dictate whether pathogen transmission rates skew towards males or females. This recognition of sex-specific plasticity not only broadens the search for evolutionary generalizations of sex biases in infection outcomes (as per Hamilton and Zuk, 1982; Zuk, 1990; Folstad and Karter, 1992; Zuk and McKean, 1996; Rolff, 2002; Zuk et al., 2009) but also implies that similar sex-specific dynamics may also extent to other environmental factors (i.e., resource availability, Gipson et al. 2022; pollution, Aulsebrook et al. 2023, Polverino et al. 2023). Building this enhanced understanding of sex-specific plasticity and shifting sex-biases into studies of pathogen transmission will be essential to improve our capacity to anticipate the trajectory of infectious diseases in a sexually dimorphic and changing world.

## Acknowledgements

We thank Kate Langwig, Roland Regoes, and Gregory Markowsky for helpful discussions on host susceptibility. This project was funded by the Australian Research Council (grant no. DP200102522 to M.D.H).

