## Supplemental Figure 1 for "The sicker sex is plastic: Thermal plasticity determines sex biases in pathogen transmission"

**Supplementary material: The sicker sex is plastic: Thermal regulation of sex biases in pathogen transmission**

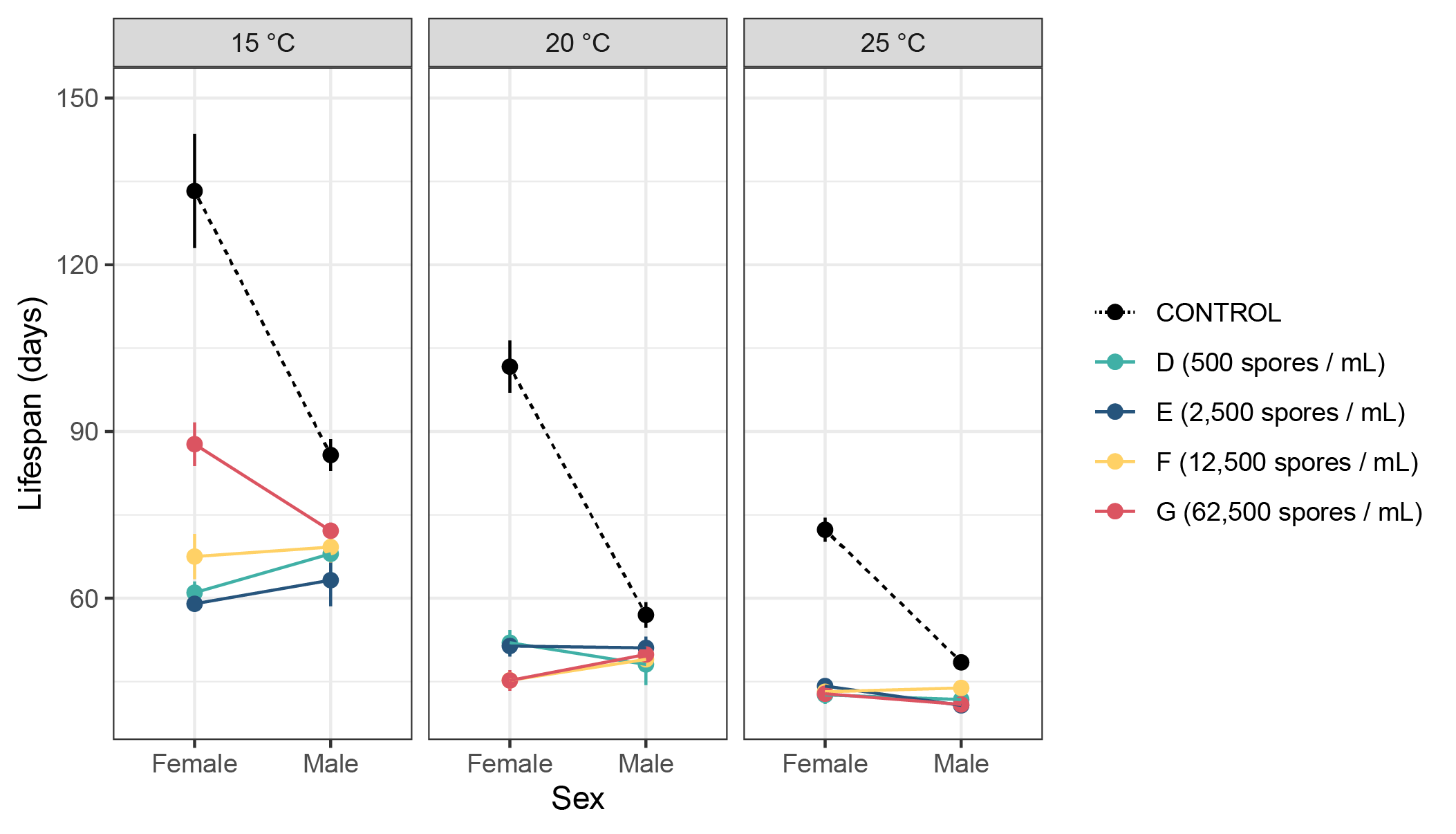

**Figure 1**. Thermal plasticity in lifespan as measured for male and female *Daphnia* *magna* (genotype: BE-OHZ-M10) exposed to the pathogen *Pasteuria ramosa* (genotype: C1). Shown are the predicted treatment means (± SE) for the change in lifespan of experimental animals exposed to 500, 2,500, 12,500, or 62,500 spores/mL relative to healthy controls (low A dose uninfected exposed).
